# Online parallel accumulation − serial fragmentation (PASEF) with a novel trapped ion mobility mass spectrometer

**DOI:** 10.1101/336743

**Authors:** Florian Meier, Andreas-David Brunner, Scarlet Koch, Heiner Koch, Markus Lubeck, Michael Krause, Niels Goedecke, Jens Decker, Thomas Kosinski, Melvin A. Park, Nicolai Bache, Ole Hoerning, Jüergen Cox, Oliver Räther, Matthias Mann

## Abstract

In bottom-up proteomics, peptides are separated by liquid chromatography with elution peak widths in the range of seconds, while mass spectra are acquired in about 100 microseconds with time-of-fight (TOF) instruments. This allows adding ion mobility as a third dimension of separation. Among several formats, trapped ion mobility spectrometry (TIMS) is attractive due to its small size, low voltage requirements and high efficiency of ion utilization. We have recently demonstrated a scan mode termed parallel accumulation – serial fragmentation (PASEF), which multiplies the sequencing speed without any loss in sensitivity (Meier *et al.*, PMID: 26538118). Here we introduce the timsTOF Pro instrument, which optimally implements online PASEF. It features an orthogonal ion path into the ion mobility device, limiting the amount of debris entering the instrument and making it very robust in daily operation. We investigate different precursor selection schemes for shotgun proteomics to optimally allocate in excess of 100 fragmentation events per second. More than 800,000 fragmentation spectra in standard 120 min LC runs are easily achievable, which can be used for near exhaustive precursor selection in complex mixtures or re-sequencing weak precursors. MaxQuant identified more than 6,400 proteins in single run HeLa analyses without matching to a library, and with high quantitative reproducibility (R > 0.97). Online PASEF achieves a remarkable sensitivity with more than 2,900 proteins identified in 30 min runs of only 10 ng HeLa digest. We also show that highly reproducible collisional cross sections can be acquired on a large scale (R > 0.99). PASEF on the timsTOF Pro is a valuable addition to the technological toolbox in proteomics, with a number of unique operating modes that are only beginning to be explored.

---

**J**ointly, proteins form a cellular machinery – the proteome - that orchestrates essentially all biological processes in health and disease. Studying it on a system-wide scale holds great promise to advance our understanding of cellular biology and disease mechanisms^1–3^. However, as compared to genomics and transcriptomics technologies, proteomics still lags behind in terms of coverage, throughput and sensitivity. Virtually complete measurements of mammalian proteomes have become possible^4^, but have mostly involved laborious sample preparation workflows, days of measurement time and substantial amounts of starting material. Furthermore, current high-performance instrumentation often requires expert knowledge and extensive maintenance, which impedes widespread adaptation of proteomics in non-specialized laboratories.

In bottom-up workflows, proteins are extracted from a biological sample of interest and enzymatically cleaved, which makes them more amenable to mass spectrometric (MS) analysis. The resulting complex peptide mixtures are typically separated via nano-flow liquid chromatography (LC), ionized by electrospray and mass analyzed. In ‘data-dependent’ or ‘topN’ acquisition schemes, the mass spectrometer detects suitable peptide precursor ions in full scans (MS) and selects them for fragmentation in *N* consecutive MS/MS scans. High resolution and high mass accuracy analyzers detect hundreds of thousands of distinct molecular features in single LC-MS experiments, of which only a minority is identified and quantified^5^. These co-eluting peptides with abundances ranging over many orders of magnitude present a formidable analytical challenge, which has constantly pushed the development of faster and more sensitive instrumentation over the last decades^1,3,6,7^.

Time-of-flight (TOF) instruments have a number of very desirable properties for the analysis of complex peptide mixtures and have consequently been employed in shotgun proteomics for a long time^8,9^. Instrumental performance has steadily improved over the years, and our groups have described shotgun proteome measurements at a resolution of more than 35,000 within about 100 μs on the *impact II*^10^, the predecessor of the instrument that is the subject of this paper. The high acquisition rate of TOF instruments allows coupling them with very fast separation techniques, such as ion mobility spectrometry^11–13^. IMS separates ions in the gas phase based on their size and shape, or more precisely their collisional cross section (CCS, Ω), typically within 10s to 100s of milliseconds^14^. As they emerge from the ion mobility device, they can be efficiently sampled in the ms or sub-ms time frame. Nested between LC and MS, the technology provides an additional dimension of separation^15–17^ and can increase analysis speed and selectivity^18^, in particular with highly complex proteomics samples^19–23^. However, many implementations of IMS, such as drift tubes, are challenging to implement due to the device sizes and high voltages involved, and may also limit the proportion of the continuous incoming beam that can be utilized^12,13,24^. Trapped ion mobility spectrometry (TIMS)^25,26^ reverses the concept of traditional drift tube ion mobility, by bringing ions to a rest at different positions in an ion tunnel device, balanced in an electrical field against a constant gas stream^27^. Once a sufficient number of ions have been trapped and separated, lowering the electrical potential releases time-resolved ions from the TIMS device into the downstream mass analyzer. This design reduces IMS analyzer dimensions to about 10 cm centimeters in length – allowing two of them to be implemented in series for 100% duty cycle operation^28^. TIMS furthermore offers high flexibility in that users can tune the ion mobility resolving power (Ω/Δ_FWHM_Ω) up to 200 or higher by simply lowering the TIMS scan speed^29,30^.

We have recently introduced ‘Parallel Accumulation – SErial Fragmentation’ (PASEF)^31^, which synchronizes MS/MS precursor selection with TIMS separation. This acquisition scheme allows fragmentation of more than one precursor per TIMS scan. We demonstrated that PASEF increases the sequencing speed several-fold without loss of sensitivity. As precursor ions are accumulated in parallel, PASEF overcomes the diminishing returns of increasingly fast MS/MS acquisition, which otherwise necessarily implied less and less ions per spectrum. Our first iteration was implemented on a laboratory prototype, which required manual precursor programming and was limited by the speed of the electronics involved. Here, we describe the construction and investigate the proteomics performance of the first mass spectrometer that fully integrates the PASEF concept, the Bruker *timsTOF Pro*.

## EXPERIMENTAL PROCEDURES

### Cell culture and sample preparation

Human cervical cancer cells (HeLa S3, ATCC, USA) were grown in Dulbecco’s modified Eagle’s medium with 10% fetal bovine serum, 20 mM glutamine and 1% penicillin-streptomycin (all PAA Laboratories, Germany). Escherichia coli (strain: XL1 blue) was cultured at 37 °C in LB medium until logarithmic phase (optical density = 0.5, λ = 600 nm). Cells were collected by centrifugation. Following a washing step with cold phosphate buffered saline, they were pelleted and flash frozen in liquid nitrogen and stored at −80 °C.

One-device cell lysis, reduction, and alkylation was performed in sodium deoxycholate (SDC) buffer with chloroacetamide (PreOmics GmbH, Germany) according to our previously published protocol^32^. Briefly, the cell suspension was twice boiled for 10 min at 95 °C and subsequently sonicated for 15 min at maximum energy (Bioruptor, Diagenode, Belgium). Proteins were enzymatically hydrolyzed overnight at 37 °C by LysC and trypsin (1:100 enzyme:protein (wt/wt) for both). To stop the digestion, the reaction mixture was acidified with five volumes of isopropanol with 1% trifluoroacetic acid (TFA). Peptides were de-salted and purified in two steps, first on styrenedivinylbenzene-reversed phase sulfonate (SDB-RPS), and second on C_18_ sorbent. The dried eluates were re-constituted in water with 2% acetonitrile (ACN) and 0.1% TFA for direct LC-MS analysis or high pH reverse-phase fractionation.

### Peptide fractionation

High pH reversed-phase fractionation was performed on an EASY-nLC 1000 (Thermo Fisher Scientific, Germany) coupled to a ‘spider fractionator’ (PreOmics GmbH, Martinsried, Germany) as detailed in ref ^33^. Purified peptides were separated on a 30 cm x 250 μm reversed-phase column (PreOmics) at a flow rate of 2 μL/min at pH 10. The binary gradient started from 3% buffer B (PreOmics), followed by linear increases to first 30% B within 45 min, to 60% B within 17 min, and finally to 95% B within 5 min. Each sample was automatically concatenated into 24 fractions in 90 s time intervals. The fractions were dried in a vacuum-centrifuge and re-constituted in water with 2% ACN and 0.1% TFA for LC-MS analysis.

### Liquid Chromatography

An EASY-nLC 1200 (Thermo Fisher Scientific, Germany) ultra-high pressure nano-flow chromatography system was coupled online to a hybrid trapped ion mobility spectrometry – quadrupole time of flight mass spectrometer (*timsTOF Pro*, Bruker Daltonics) with a modified nano-electrospray ion source^10^ (CaptiveSpray, Bruker Daltonics). Liquid chromatography was performed at 60 °C and with a constant flow of 400 nL/min on a reversed-phase column (50 cm x 75 μm i.d.) with a pulled emitter tip, packed with 1.9 μm C_18_-coated porous silica beads (Dr. Maisch, Germany). Mobile phases A and B were water with 0.1% formic acid (vol/vol) and 80/20/0.1% ACN/water/formic acid (vol/vol/vol), respectively. In 120 min experiments, peptides were separated with a linear gradient from 7.5 to 27.5% B within 60 min, followed by an increase to 37.5% B within 30 min and further to 55% within 10 min, followed by a washing step at 95% B and re-equilibration. In 60 min separations, the gradient increased from 10 to 30% B within 30 min, followed by an increase to 40% B within 15 min and further to 57.5% B within 5 min before washing and re-equilibration. In 30 min separations, the initial 10-30% B step was 15 min, followed by a linear increase to 40% B (7.5 min) and 57.5% B (2.5 min) before washing and re-equilibration.

For some experiments we used the Evosep One (Evosep, Odense, Denmark), a new HPLC instrument employing an embedded gradient and capable of fast turnaround between analyses^34^. Samples were eluted from Evotips at low pressure into the storage loop with a gradient offset to lower the percentage of organic buffer. Separation was performed on a customized 5.6 min gradient (200 samples/day method) at a flow rate of 1.5 μL/min on a 4 cm x 150 μm i.d. reversed-phase column packed with 3 μm C_18_-coated porous silica beads (PepSep, Odense, Denmark).

### The timsTOF Pro mass spectrometer

The *timsTOF Pro* is the successor to the *impact II* instrument, compared to which it features an additional ion mobility region. However, the *timsTOF Pro* is a complete redesign in hardware and firmware. Apart from incorporating TIMS, the design goals included the achievement of similar or better mass resolution (>35,000) and improved robustness through a changed ion path.

In the experiments described here, the mass spectrometer was operated in PASEF mode. Desolvated ions entered the vacuum region through the glass capillary and were deflected by 90°, focused in an electrodynamic funnel, and trapped in the front region of the TIMS tunnel consisting of stacked printed circuit boards (PCBs) with an inner diameter of 8 mm and a total length of 100 mm. The PCB electrodes form a stacked multipole in the direction of ion transfer. An applied RF potential of 350 V_*pp*_ confined the trapped ions radially. The TIMS tunnel is electrically separated into two parts (‘dual TIMS’), where the first region is operated as an ion accumulation trap that stores and pre-separates ions according to their mobility, and the second part performs trapped ion mobility analysis in parallel. Note that equal accumulation and analysis times in both TIMS regions enable operation at duty cycles up to 100%. Ion transfer between the two regions takes 2 ms and therefore does not affect the overall ion utilization for typical ramp and accumulation times around 50 to 200 ms.

In both TIMS regions, the RF field is superimposed (from entrance to exit) by an increasing longitudinal electrical field gradient, such that ions in the tunnel simultaneously experience a drag from the incoming gas flow through the capillary and a repulsion from the electrical field. Depending on their collisional cross sections and charge states, they come to rest closer to the entrance of the tunnel (high ion mobility) or closer to its exit (low ion mobility). Trapped ion mobility separation was achieved by ramping the entrance potential of the second TIMS region from −207 V to −77 V. A single TIMS-MS scan is composed of many individual TOF scans of about 110 μs each. In the experiments reported here, we systematically varied the ramp times from 50, 100, 150, to 200 ms while keeping the duty cycle fixed at 100%. The quantification benchmark experiment, the 60 min dilution series and the high pH reverse-phase fractions were each acquired with a 100 ms ramp and 10 PASEF MS/MS scans per topN acquisition cycle; the 30 min dilution series was acquired with a 50 ms ramp and 10 PASEF MS/MS scans per cycle; experiments on the Evosep One were performed with a 100 ms ramp and four PASEF MS/MS scans per cycle.

MS and MS/MS spectra were recorded from *m/z* 100 to 1,700. Suitable precursor ions for PASEF-MS/MS were selected in real time from TIMS-MS survey scans by a sophisticated PASEF scheduling algorithm (see also Results). A polygon filter was applied to the *m/z* and ion mobility plane to select features most likely representing peptide precursors rather than singly charged background ions. The quadrupole isolation width was set to 2 Th for m/z < 700 and 3 Th for m/z > 700, and the collision energy was ramped stepwise as a function of increasing ion mobility: 52 eV for 0-19% of the ramp time; 47 eV from 19-38%; 42 eV from 38-57%; 37 eV from 57-76%; and 32 eV for the remainder.

The TIMS elution voltage was calibrated linearly to obtain reduced ion mobility coefficients (1/K_0_) using three selected ions of the Agilent ESI-L Tuning Mix (*m/z* 622, 922, 1222)^35^.

Collisional cross sections were calculated from the Mason Schamp equation^36^:

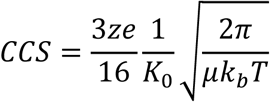

where *z* is the charge of the ion, *e* is the elemental charge, *k*_*b*_ is Boltzman’s constant, *μ* is the reduced mass, and *T* the temperature (305 K).

### Data analysis

Mass spectrometry raw files were processed with MaxQuant^37^ version 1.6.1.12, which has been extended to incorporate the additional ion mobility dimension and adapted to handle the TIMS data format. This new version of MaxQuant is publicly available and will be described in detail separately (Cox and co-workers, *in preparation*). Briefly, it assembles four-dimensional isotope clusters-defined by *m/z*, retention time, ion mobility and intensity - from the TIMS-MS spectra and extracts ion mobility separated MS/MS spectra from the PASEF scans. Each MS/MS spectrum is assigned to its respective precursor ions by quadrupole isolation *m/z* and ion mobility values, and in case a precursor has been fragmented multiple times in one acquisition cycle, the respective spectra are collapsed to a single spectrum with increased signal-to-noise. The ‘TIMS half width’ parameter was set to 4 TOF triggers, the ‘TIMS step width’ to 3, the ‘TIMS mass resolution’ to 32,000 and MS/MS peaks with an intensity below 1.5 units were discarded.

The MS/MS spectra were matched to *in silico* derived fragment mass values of tryptic peptides from a reference proteome (Uniprot, 2016/05, HeLa: 91,618 entries including isoforms, E.coli: 4,313 entries including isoforms) and 245 potential contaminants by the built-in Andromeda search engine^38^. A maximum of two missing cleavages were allowed, the required minimum peptide sequence length was 7 amino acids, and the peptide mass was limited to a maximum of 4,600 Da. Carbamidomethylation of cysteine residues was set as a fixed modification, and methionine oxidation and acetylation of protein N-termini as variable modifications. The initial maximum mass tolerances were 70 ppm for precursor ions and 35 ppm for fragment ions. We employed a reversed sequence library to control the false discovery rate (FDR) at less than 1% for peptide spectrum matches and protein group identifications.

Decoy database hits, proteins identified as potential contaminants, and proteins identified exclusively by one site modification were excluded from further analysis. Label-free protein quantification was performed with the MaxLFQ algorithm^39^ requiring a minimum ratio count of 1. All other MaxQuant parameters were kept at their default values.

Mass spectrometric metadata, such as the information about PASEF-scheduled precursor ions, were directly accessed and extracted from the Bruker *.tdf* raw files with a SQLite database viewer (SQLite Manager, v0.8.3.1). Bioinformatic analysis and visualization was performed in either Python (Jupyter Notebook), Perseus^40^ (v1.6.0.8) or the R statistical computing environment^41^ (v3.2.1).

### Experimental Design and Statistical Rationale

Samples were grouped by mass spectrometric acquisition methods or, in case of the data for Fig. 5, by pipetting ratios. Replicate injections were performed to assess the technical reproducibility of the respective methods and their quantitative accuracy. To allow accurate external calibration of ion mobility values, we acquired experiments with different TIMS ramp times in batches. Dilution series were measured from low to high concentrations starting with blank runs to avoid carry over. This study does not draw biological conclusions, which is why process and biological replicates or controls were not performed.

## RESULTS

### Construction of a TIMS-QTOF instrument with online PASEF

The *timsTOF Pro* is a quadrupole time-of-flight (QTOF) mass spectrometer equipped with a second generation dual TIMS analyzer in the first vacuum stage (**Fig. 1**). This set-up spatially separates ion accumulation and ion mobility analysis into two sequential sections of the TIMS tunnel, so that these steps happen in parallel^28^ (analyzer 1 and 2 in **Fig. 1b**). Within the limits of ion storage capacity, up to 100% of the ions that enter the mass spectrometer can therefore be utilized for mass analysis. Here, we typically accumulated ions for 50 to 200 ms, and transferred them into the second TIMS region within 2 ms. From this TIMS region they were released by decreasing the voltage gradient in a linear manner within 50 to 200 ms (TIMS ‘ramp time’). Simulations show that most of the ion mobility separation happens near the top plateau close to the exit of the device^42–44^ and we observed that leaving peptide ion packets had narrow ion mobility peaks with median half widths of about 2 ms or less (**Fig. 1c**). In TIMS, low mobility ions are released or ‘eluted’ first, followed by more mobile ions with smaller collisional cross sections relative to their charge. In addition to separating ions by shape and size, the time-focusing effect of TIMS increases signal-to-noise ratios about 50-fold (depending on the relative accumulation and ramp times) compared with the standard continuous acquisition mode because ion species are concentrated into narrow packets whereas the noise distributes across the ion mobility scan^28^.

**Figure 1.**
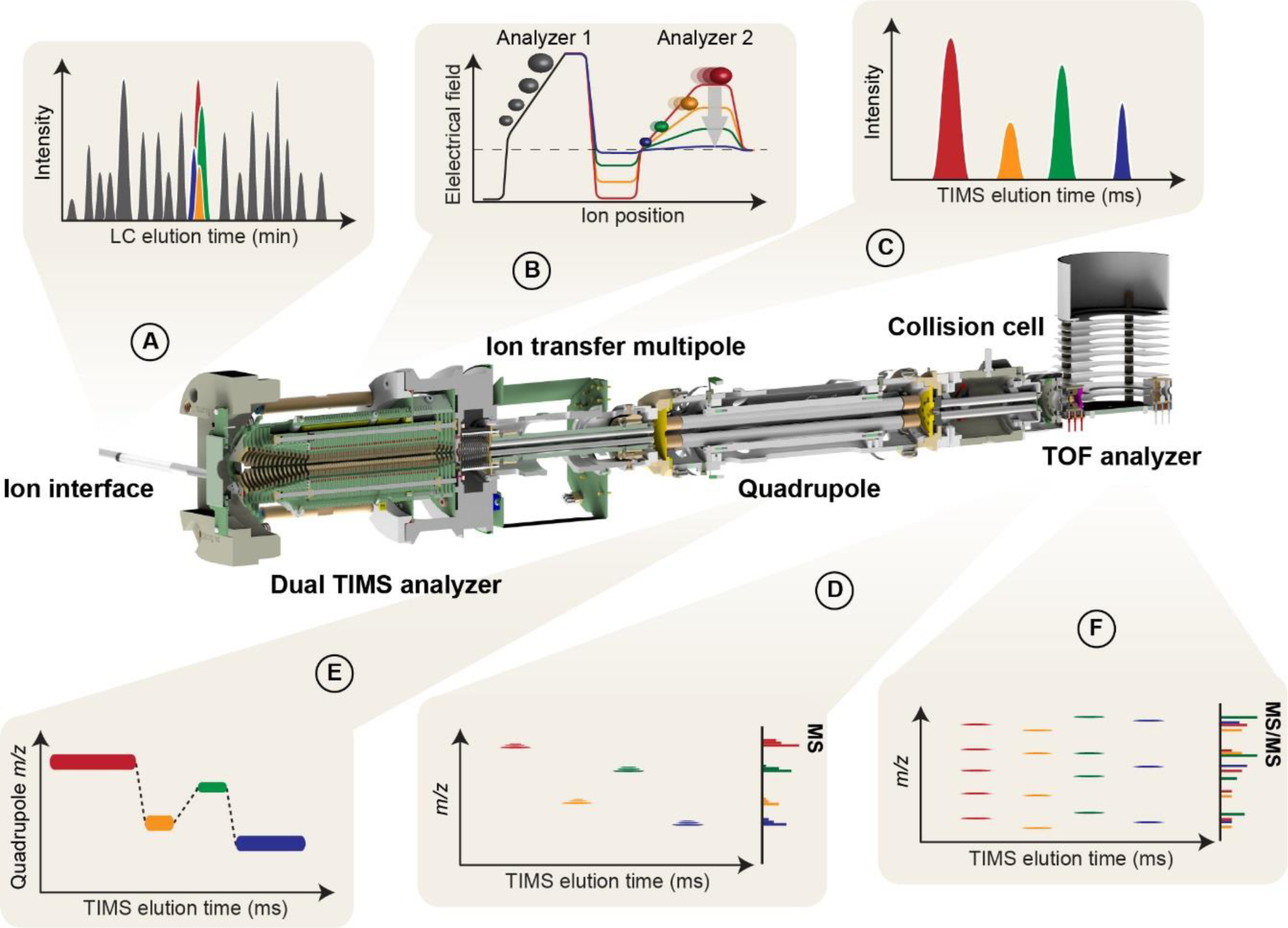
Online Parallel Accumulation - Serial Fragmentation (PASEF) with the timsTOF Pro. (**A**) Peptides eluting from the chromatographic column are ionized and enter the mass spectrometer through a glass capillary. (**B**) In the dual TIMS analyzer, the first TIMS section traps and stores ion packets, and the second resolves them by mobility. (**C, D**) Ion mobility separated ions are released sequentially from the second TIMS analyzer as a function of decreasing electrical field strength and yield mobility-resolved mass spectra. (**E**) In PASEF MS/MS scans, the TIMS analyzer and the quadrupole are synchronized and the quadrupole isolation window switches within sub-milliseconds between mobility resolved precursor ions of different *m/z*. (**F**) This yields multiple ion mobility resolved MS/MS spectra from a single TIMS scan, and ensures that multiple trapped precursor ion species are used for fragmentation. Non mobility-resolved MS and MS/MS spectra are projected onto the right axes in D and F for comparison.

At the exit of the TIMS device, ions pass through the ion transfer multipole and the quadrupole mass filter and are accelerated into the collision cell. From there, intact (MS scans) or fragment (MS/MS scans) ions are extracted into an orthogonal accelerator unit and pushed into the flight tube for mass analysis (**Fig. 1d**). The ions enter a V-shaped flight path through a two-stage reflectron and finally impinge on a multi-channel plate (MCP) ion detector coupled to a 10-bit digitizer with a sampling rate of 5 Gigasamples(GS)/s, enabling high-resolution mass analysis (R > 35,000 throughout the entire mass range). We observed that the re-designed ion transfer path – presumably mainly the 90 degree bent at the entrance of the TIMS device and the new quadrupole with increased inner diameter - had a positive effect on the robustness. This was evidenced by continuous operation of the instrument during its development for more than 1.5 years, in which time we only cleaned the ion transfer capillary but not the internals of the instrument.

In PASEF mode, MS/MS precursor selection by the quadrupole mass filter is synchronized with the release of ions from the TIMS device, which requires very fast switching times of the quadrupole to keep pace with the fast ion mobility separation and to maximize the number of precursors per TIMS scan (**Fig. 1e**). The *timsTOF Pro* electronics have been designed to meet these requirements and RF and DC voltages for mass selection are now calculated and set by a real-time field-programmable array (FPGA), as opposed to a conventional and slower serial interface. This allows fully synchronized operation of TIMS and quadrupole with switching times of 1 ms or less. By setting the quadrupole to *N* different *m/z* windows, PASEF yields *N* ion-mobility-resolved MS/MS spectra for a single TIMS scan (**Fig. 1f**). Because all precursor ions are stored in parallel, the absolute ion count per MS/MS spectrum is equal to a conventional TOF MS/MS spectrum summed up over the accumulation time, giving rise to an *N*-fold increase in sequencing speed without sacrificing sensitivity. The maximum number of precursors per TIMS scan is not limited by the instrument electronics, but rather by the separation of precursors in the ion mobility dimension and by the efficient design of ‘switching routes’ for precursor selection, which will be described next.

### PASEF precursor selection in real-time

In complex proteomics samples, such as whole cell lysates, hundreds to thousands of peptides elute at any time, presenting a challenge for optimal selection even with the ten-fold higher sequencing speed offered by PASEF. Fortunately, precursors are now distributed in a two-dimensional (*m/z* and ion mobility) space in which an optimal route can be selected, similar to the ‘travelling salesman problem’ in computer science. Even though exact solutions exist, for example by a brute-force method that simply iterates over all possible combinations, they cannot be computed on the LC time scale nor is it clear which peaks are most desirable to ‘visit’. Instead, we here developed a heuristic algorithm that limits the computational time to about 100 ms in complex samples, and aims to maximize the number of precursors per acquisition cycle that can be successfully identified. This involves three dimensions – precursor *m/z*, signal intensity and ion mobility (**Fig. 2**). Our precursor search is offset by one acquisition cycle from ongoing data acquisition to avoid introducing any scan overhead time. In distributing precursors to PASEF scans, our algorithm accounts for the quadrupole switching time as well as the elution order of ion mobility peaks and prioritizes high-abundance precursors. In principle, the maximum coverage of eluting peptides should be achieved by using the PASEF speed advantage exclusively on unique precursor ions. However, this leads to many low abundant precursors being selected, and thus many low-quality MS/MS spectra. An alternative strategy is to deliberately re-sequence selected low-abundance precursor ions in subsequent PASEF scans to obtain summed spectra with increased signal-to-noise. This is implemented in our precursor algorithm by a ‘target intensity’ parameter, with which users can balance the desired spectral quality with the number of unique precursors. Other than that, we excluded precursors dynamically after one sequencing event to not compromise proteomic depth. Singly-charged species were readily excluded by their characteristic positions in the *m/z* vs. ion mobility plane. The flow chart in **Supplementary Fig. 1** depicts the precursor selection algorithm in detail.

**Figure 2.**
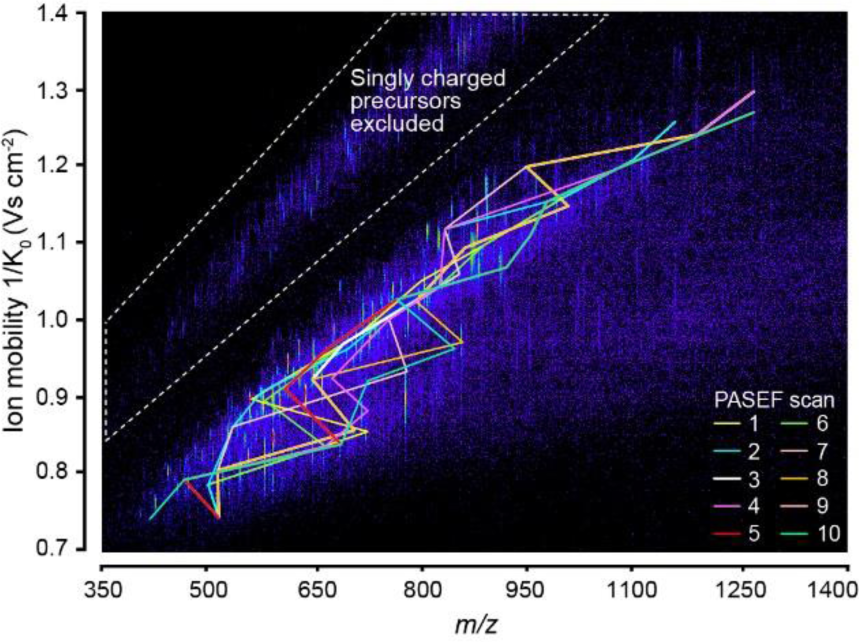
Real-time PASEF precursor selection in three dimensions. Heat-map visualization of ion mobility resolved peptide ions at a single time point in an LC-TIMS-MS analysis of a HeLa digest. Connected lines indicate the *m/z* and mobility positions of all precursor ions selected for fragmentation in the following TIMS-PASEF scans (color-coded).

We tested the performance of our precursor selection algorithm in 120 min LC-TIMS-MS runs of HeLa digests. **Figure 2** shows a representative TIMS-MS survey scan in the middle of the LC gradient. From this 100 ms TIMS scan, our algorithm selected 50 unique precursor ions for fragmentation in the subsequent PASEF scans (color-coded) out of which 32 low-abundance precursors were repeatedly sequenced. All precursor ions were widely distributed in *m/z* and ion mobility space, indicating an efficient coverage of the entire precursor space. In total, 118 MS/MS spectra were acquired in this cycle, which equals a sequencing rate of more than 100 Hz. Because all precursors were accumulated for 100 ms, the total number of ions for each precursor corresponds to that of a 10 Hz MS/MS selection if no PASEF had been employed.

With the selection algorithm in place, we inspected hundreds of precursor identifications in our data sets. Often, the separation of precursors along the additional ion mobility dimension was crucial as illustrated in **Figure 3**. In a projection of the data onto the *m/z* axis, no obvious precursor signals were present, even when enlarging the signal ten-fold relatively to the more abundant peaks. However, the precursor selection algorithm had found and fragmented two distinct isotope clusters in ion mobility – *m/z* space, which were separately fragmented by PASEF and clearly identified (**Supplementary Fig. 2**).

**Figure 3.**
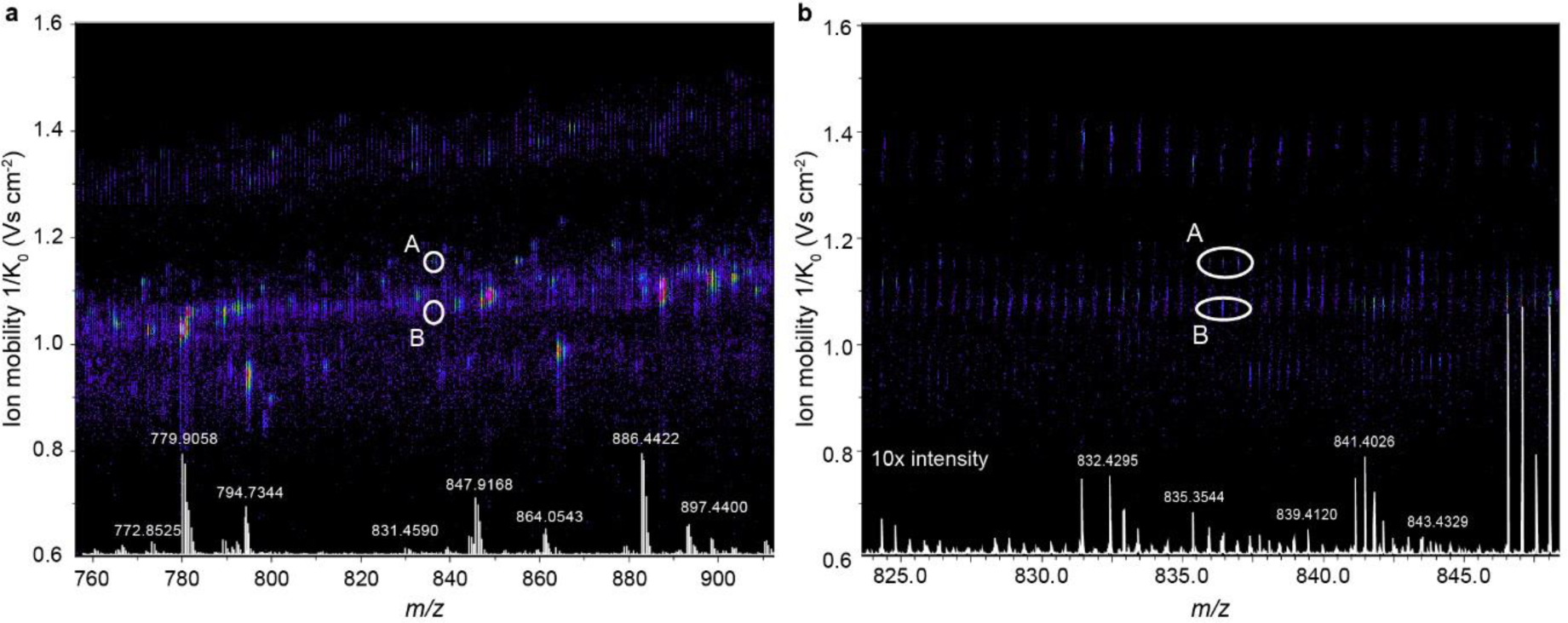
Trapped ion mobility separation of peptide precursorions. (**a**) The two nearly isobaric peptide ions A and B were distinguished by their ion mobility and selected separately for fragmentation by the PASEF scheduling algorithm in an LC-TIMS-MS experiment of a HeLa digest. (**b**) Zoomed view into the precursor *m/z* range. Non mobility-resolved MS spectra are projected onto the lower axis for comparison. The corresponding MS/MS spectra are shown in Suppl. Fig. 1.

### Single run proteomes

Next, we investigated the effect of different TIMS ramp times on precursor selection. Given a minimum selection and transition time for the quadrupole adjustment of a few ms, the overall number of achievable fragmentation events should be roughly similar for different TIMS ramp times as increasing ramp time allows fragmenting more precursors per PASEF scan-while acquiring less scans overall. To find a good balance for proteomics applications, we varied the TIMS ramp from 50 to 200 ms and kept the PASEF scans at 10 per acquisition cycle. We chose to operate the instrument at a near 100% duty cycle by setting the TIMS acquisition time equal to the ramp time.

With the slowest (and therefore highest mobility resolving) TIMS ramp, an average of 23.3 precursors were sequenced per scan (**Fig. 4a**). Faster ramp times resulted in nearly proportionately less precursors per PASEF scan, but due to the higher number of scans per analysis, faster scans generated more MS/MS events in total - up to a remarkable 840,000 spectra in two hours (**Fig. 4b**). For comparison, acquiring the same number of MS/MS spectra without PASEF at the same sensitivity would have taken 12 times longer - about one day. For all ramp times, the instrument was sequencing at rates above 100 Hz during the time that peptides were eluting. We decided to use this extreme speed in part on re-sequencing low-abundance peptides to generate higher-quality summed spectra (**Fig. 4c**). On average, a given precursor ion was fragmented 2.1 times in 50 ms ramps and 3.1 times with 200 ms ramps. Overall, this resulted in up to 380,000 MS/MS spectra of unique precursor ions in a single run as detected by the real-time PASEF scheduling algorithm, although post-processing in MaxQuant combined many of these (**Fig. 4d**).

**Figure 4.**
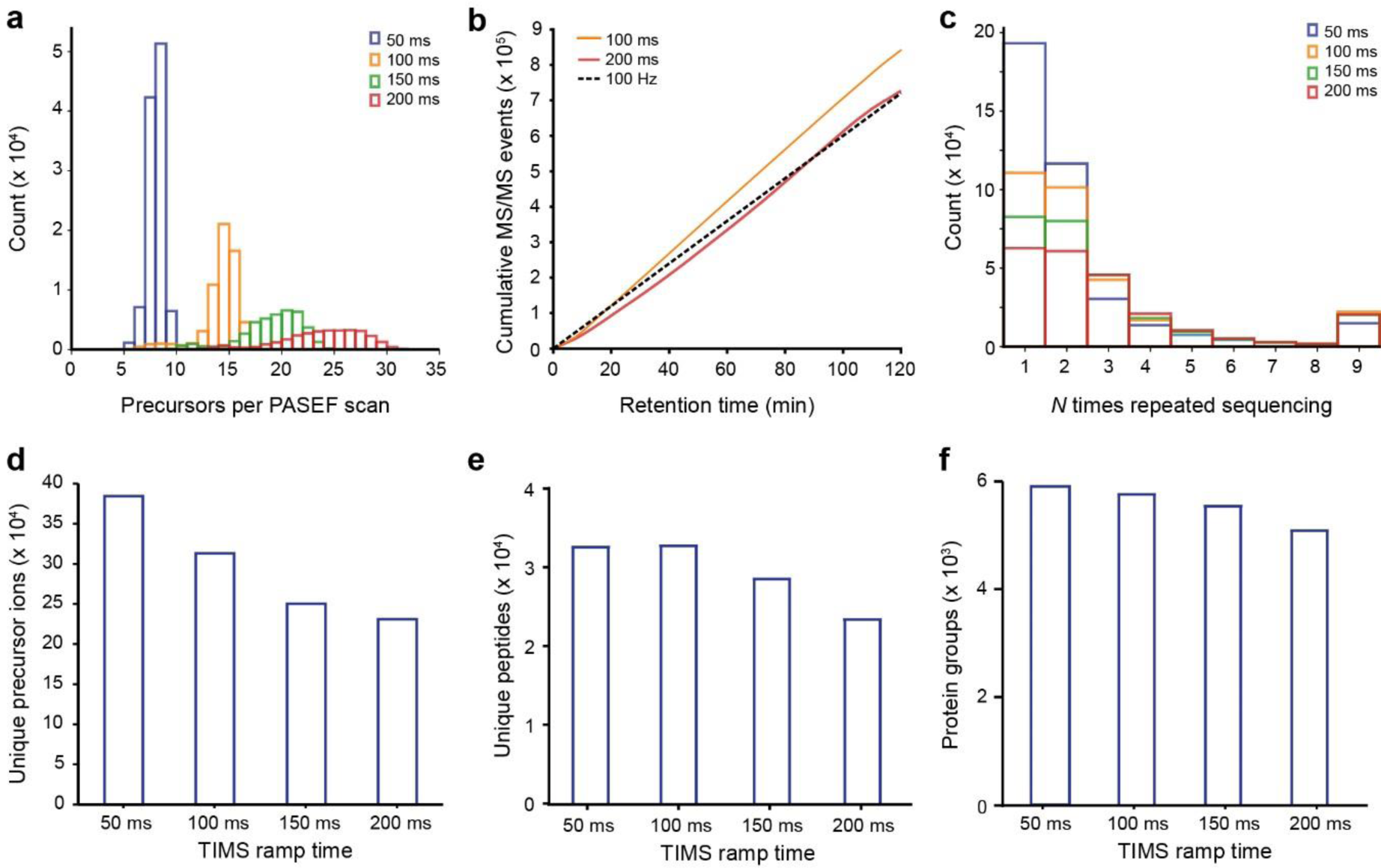
Single run analyses of a HeLa digest. (**a**) Number of selected precursor ions per PASEF scan with different TIMS ramp times in 120 min runs of 200 ng HeLa digests. (**b**) Cumulative number of PASEF MS/MS spectra as a function of retention time for 100 ms and 200 ms TIMS ramps. The dashed line indicates the theoretical number of MS/MS spectra for a constant acquisition rate of 100 Hz (**c**) Number of repeated sequencing events for precursors with different ramp times. (**d**) Number of unique precursor ions detected with different TIMS settings. (**e**) Average number of sequence-unique peptides identified in a single run (N=4) with different TIMS settings. (**f**) Average number of protein group identifications in a single run (N=4) with different TIMS settings.

From 200 ng whole-cell HeLa digest per run, we identified on average 23,696 sequence-unique peptides in quadruplicate single runs with the 200 ms method, and about 33,000 with the faster 50 ms and 100 ms methods (**Fig. 4e**). Average peptide length was 15 amino acids, similar to that expected from in silico digests of the UniProt database given our minimum peptide length of seven. The number of inferred protein groups at a false discovery rate (FDR) below 1% increased to an average of 5,970 protein groups per run with decreasing TIMS ramp times from 200 to 50 ms (**Fig. 4f**). With the 50 ms ramps, we identified in total 6,491 protein groups (5,753 with two or more peptides) with a median sequence coverage of 19.9%. This is an excellent value given the very low starting amount and the absence of fractionation or a matching library.

**Figure 5.**
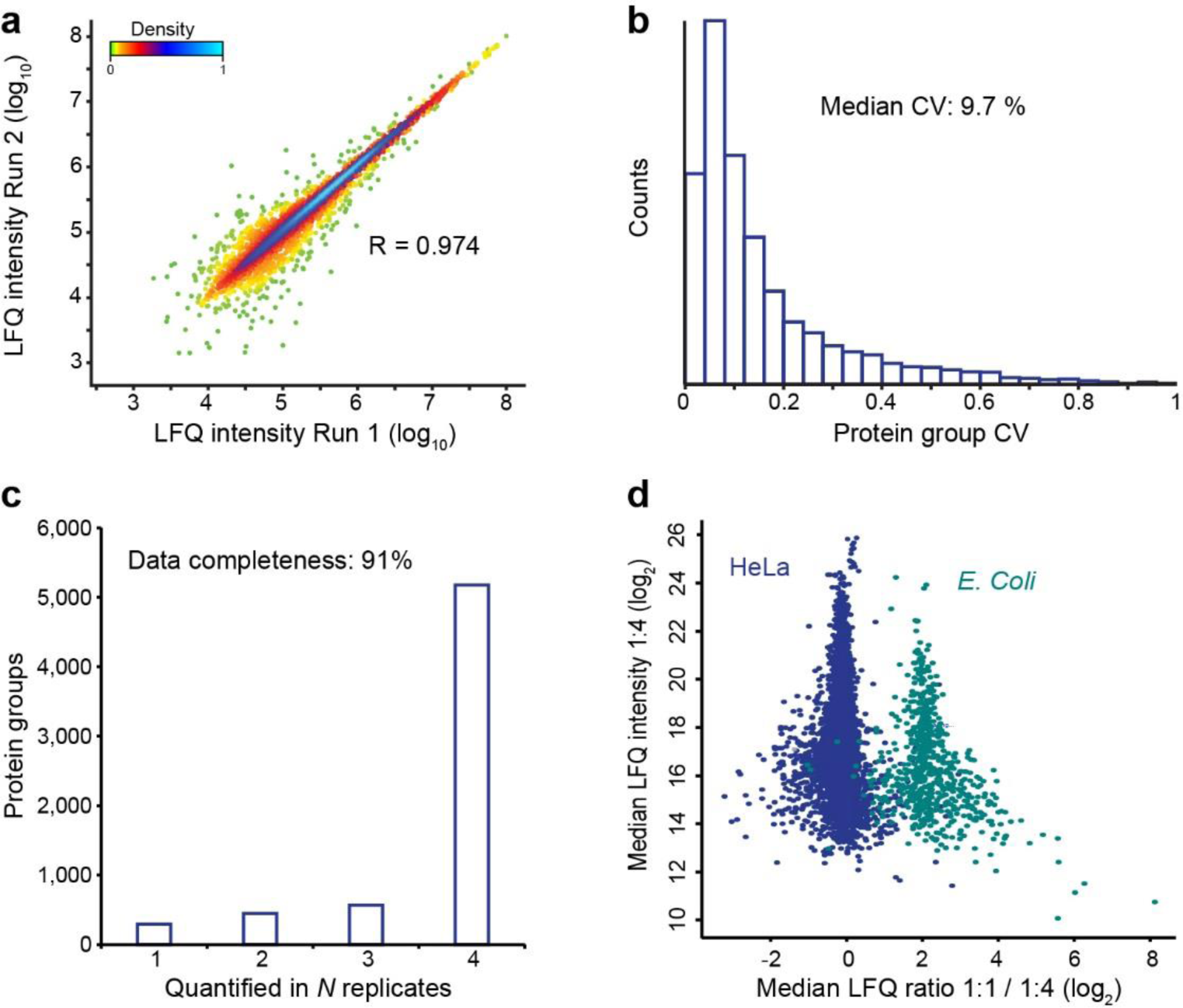
Label-free proteome quantification. (**a**) Pearson correlation of protein intensities in two replicate injections of a HeLa digest. (**b**) Coefficients of variation (CVs) for protein quantities in four replicates. (**c**) Number of proteins quantified in *N* out of four replicates. (**d**) Label-free quantification benchmark with whole-cell HeLa and E.coli digests mixed in 1:1 and 1:4 ratios (wt:wt). The scatterplot shows the median fold-change of human and E.coli proteins in quintuplicate single runs.

### Label-free proteome quantification

A central task in proteomics is the accurate quantification of protein abundances across multiple biological samples. Label-free quantification (LFQ) is a popular method for this due to its simplicity, and it scales well to larger sample cohorts. Using the optimized 50 ms TIMS method we quantified on average 5,903 protein groups in 2 h LC-MS time across quadruplicate injections. Run-to-run reproducibility was high with a median pairwise Pearson correlation coefficient of 0.97 between the four runs, with excellent linearity over 4.5 orders of magnitude in protein abundance (**Fig. 5a**). The median coefficients of variation were 19.7% for the non-normalized peptide intensities and 9.7% at the protein level after MaxLFQ normalization^39^ (**Fig. 5b**).

Quantitative accuracy in proteomics may be limited if proteins are inconsistently measured across the samples. In data-dependent acquisition schemes, this is partially due to semi-stochastic precursor selection – a consequence of the large number of co-eluting precursor candidates and the finite sequencing speed. We asked if the several-fold faster PASEF method as compared with standard shotgun acquisition methods would improve this situation even without transferring identifications by precursor mass (‘matching between runs’). Indeed, PASEF alleviated the ‘missing value’ problem and provided quantification values for 5,177 proteins in four out of four runs (**Fig. 5c**). Only 294 low-abundance proteins were exclusively quantified in a single replicate. This translated into a data completeness of 91%, which compares favorably to standard data-dependent acquisition and is similar to data-independent acquisition schemes. We expect that transferring identifications between runs, as with the MaxQuant ‘matching between run’ feature, will lead to even more consistent protein quantification across samples.

To further benchmark the quantitative accuracy of our setup, we mixed tryptic digests from HeLa and *Escherichia coli* in 1:1 and 1:4 ratios and measured each sample in quintuplicate 120 min single runs. This quantified 5,268 protein groups (4,565 HeLa; 703 E.coli) in at least one out of five replicates in both experimental conditions. Plotting the median fold-changes yielded two distinct clouds for HeLa and *E.coli* proteins, which were 4.6-fold separated in abundance, slightly more than the intended 4-fold mixing ratio (**Fig. 5d**). Both populations were relatively narrow (σ(HeLa) = 0.44; σ(*E.coli*) = 0.81) and they had minimal overlap. Without imputation, a one-sided Student’s t-test returned 588 significantly changing *E.coli* proteins with at least two valid values in each group (of 621) at a permutation-based FDR below 0.05. This represents an excellent sensitivity of ∼95% and at the same time, only 64 human proteins (1.5%) were false classified as changing. From these results, we conclude that the combination of TIMS and PASEF provides precise and accurate label-free protein quantification at a high level of data completeness.

### High throughput and limited sample amounts

The performance characteristics discussed so far suggest that the instrument is particularly well suited for rapid and high sensitivity proteome analysis. To test this, we first reduced the peptide amount on column from 100 ng down to 10 ng HeLa digest per injection (**Fig. 6a**). With 100 ng on column and a 1 h gradient, we reproducibly identified 4,515 protein groups, 76% of the proteome coverage with 200 ng in half the measurement time (**Fig. 6b**). Out of these, 3,346 protein groups were quantified with a CV below 20%. At 50 ng, we identified over 4,000 protein groups with high quantitative accuracy (median CV 9.8%), motivating us to inject even lower sample amounts. Remarkably, from only 10 ng HeLa digest, we still identified 2,741 protein groups on average and 3,160 in total (2,322 with two or more peptides in at least one replicate). Assuming 150 pg protein per cell^45^, this corresponds to the total protein amount of only about 60 HeLa cells), suggesting that TIMS-PASEF is well suited to ultrasensitive applications in proteomics. Even at this miniscule sample amount, quantitative accuracy remained high with a median peptide intensity CV of 9.2% and 1,890 proteins quantified at a CV < 20%.

To investigate achievable throughput, we repeated our sensitivity experiments with a 30 min gradient (**Fig. 6c,d**). Because of the very high sequencing speed of PASEF, reducing the measurement time had only limited effect on proteome coverage. From 100 ng HeLa digest we identified on average 3,649 protein groups in quadruplicate single runs, whereas 10 ng yielded 2,536 protein groups, all with median CVs below 12%. For the 10 ng runs, this represents 93% of the proteome coverage of the 60 min single runs in half the time.

At the very short gradients made possible by the PASEF principle, throughput starts to be severely affected by the washing, loading and equilibration steps of the HPLC between injections. We therefore turned to the recently introduced Evosep One instrument, which features a preformed gradient, increasing robustness and largely eliminating idle time between injections^34^. To explore the throughput limits of complex proteome analysis with PASEF, we made use of the ‘200 samples/day’ method on the Evosep One, which consists of a 5.6 min gradient with 7.2 min total time between injections. Remarkably, in ten replicates, more than 1,100 proteins were identified on average without any identification transfer from libraries and with only 50 ng of injected cell lysate (**Fig. 6e,f**). This combination of fast LC turnaround times with PASEF also holds great promise for rapid yet comprehensive analyses of less complex samples, for example protein interactomes, or the quantification of trace-level host cell proteins (HCPs) in recombinant biotherapeutics.

**Figure 6.**
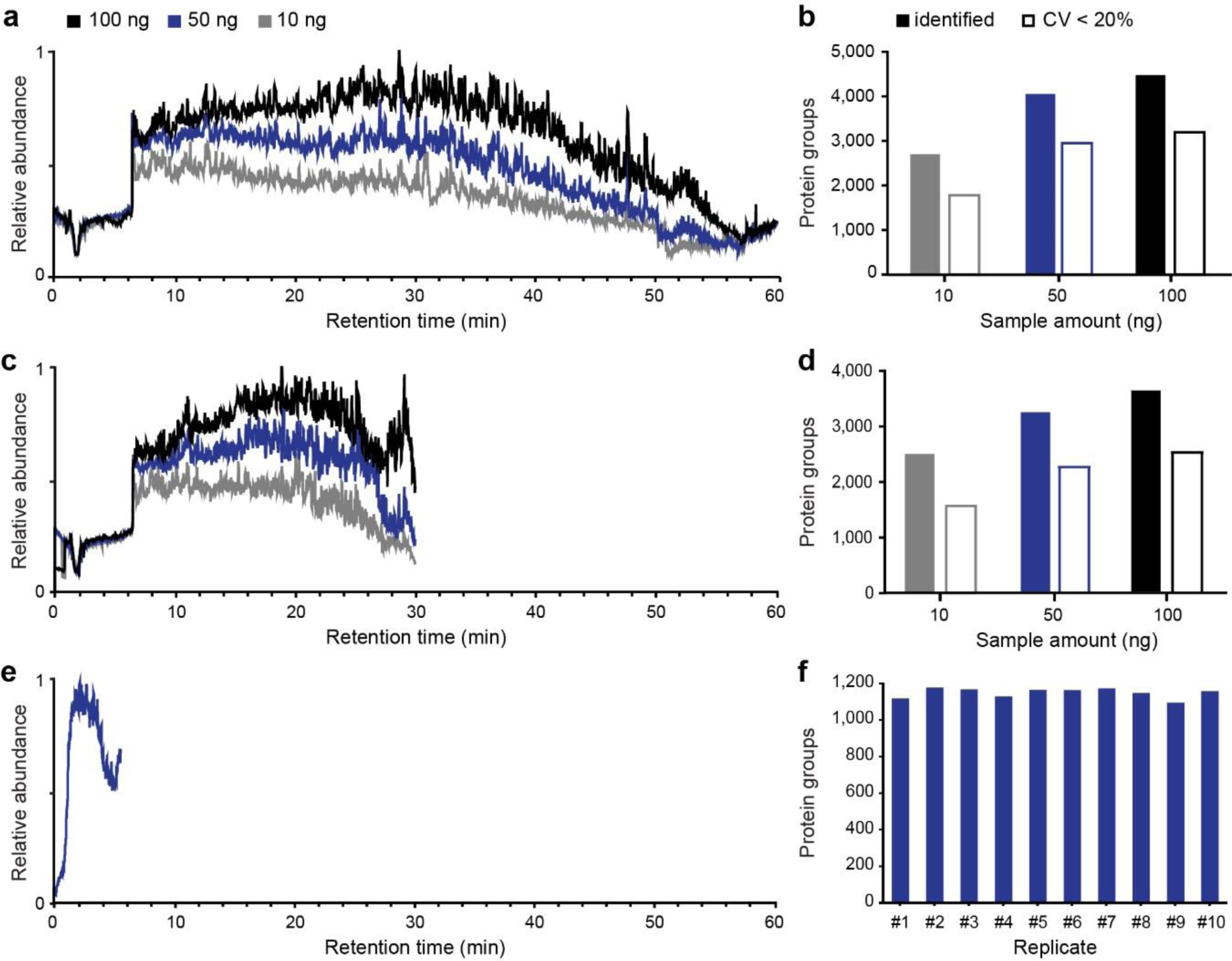
Rapid and sensitive HeLa proteome measurements. (**a**) Total ion chromatograms of the 60 min gradient and three different sample amounts on column. (**b**) Average number of protein groups identified and quantified with a CV <20% in 60 min single runs (N=3). (**c**) Total ion chromatograms of the 30 min gradient and three different sample amounts on column. (**d**) Average number of protein groups identified and quantified with a CV <20% in 30 min single runs (N=3). (**e**) Total ion chromatogram of a 5.6 min gradient with 50 ng HeLa digest on column. (**f**) Number of protein groups identified in ten replicate injections with the 5.6 min gradient.

### Large-scale measurement of peptide collisional cross sections

In TIMS, the counteracting forces of a gas flow and an electrical field are used to separate the ions and to measure their mobility. Conceptually, this closely resembles the (inverted) situation in drift tube ion mobility, where ions are dragged by an electrical field through resting gas molecules. Since the underlying physics is identical, TIMS measurements are expected to correlate directly with classical drift tube ion mobility measurements and this has been established experimentally by Park and colleagues^42^. Therefore, in contrast to other ion mobility setups^24^, such as travelling-wave ion mobility^46^ and differential ion mobility^47^, TIMS can directly determine collisional cross sections by internal or external calibration.

**Figure 7.**
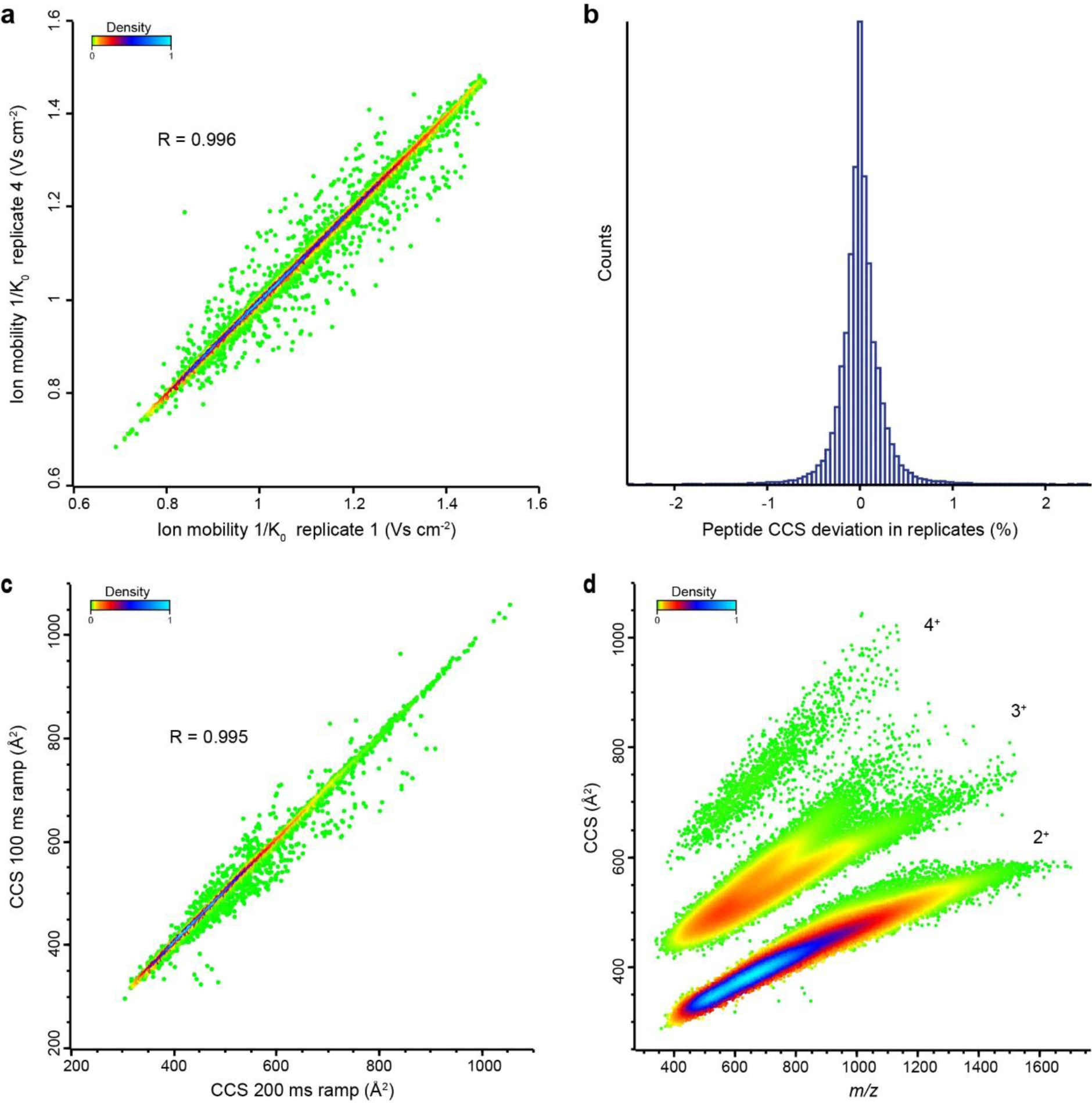
Large-scale and high-precision CCS measurements. (**a**) Pearson correlation of peptide ion mobilities in two replicate injections of a HeLa digest (100 ms TIMS ramps). (**b**) Relative deviations of CCS values of all individual peptides from their mean of quadruplicate LC-MS runs. (**c**) Pearson correlation of measured CCS values in two injections of a HeLa digest with different TIMS ramp times (100 and 200 ms TIMS ramps). (**d**) Density distribution of over 100,000 CCS values from human tryptic peptide ions as a function of *m/z*. The main populations are annotated with their respective charge states.

We reasoned that the rapid measurement of tens of thousands of peptides demonstrated above, in combination with accurate CCS measurements, should allow generating a large-scale overview of the CCS dimension of peptides. We first explored the reproducibility with repeated injections of HeLa digest. Before the first injection, we calibrated the ion mobility dimension using reduced ion mobility values (1/K_0_; Vs cm^−2^) of phosphazine derivatives from the literature^35^, which can be converted to CCS values using the Mason-Schamp equation (**Experimental Procedures**). Peptide ions can occur in multiple conformations (e.g. proline-containing peptides^48^), which results in multiple ion mobility peaks and complicates the analysis. For simplicity, we here only considered the most abundant feature reported by MaxQuant.

In four replicates, we generated 21,673 1/K_0_ values of commonly identified peptide ions in all runs with a median CV much smaller than 1% and a median pairwise correlation coefficient > 0.99 (**Fig. 7a**). Slight alterations in the gas flow can cause linear shifts in the measured mobility measurements. These can be readily taken into account by aligning the median values of all runs to the first replicate, which resulted in a median 0.1% absolute deviation of CCS values across replicates (**Fig. 7b**). In our hands, this is at least 10-fold more reproducible than LC retention time, even on the same column and with the same gradient. Interestingly, the CCS measurements were also highly transferable across different TIMS ramp times (50 ms and 150 ms) as evident from a Pearson correlation coefficient of > 0.99 between them (**Fig. 7c**).

Having established precise CCS measurements in single runs, we next used loss-less high pH fractionation^33^ to extend the scale of our data set. Measuring 24 fractions with 2 h gradients each resulted in 113,478 CCS values from 89,939 unique peptide sequences and about 9,000 protein groups. In the m/z vs. CCS plot, doubly, triply and higher-charged populations are clearly separated (**Fig. 7d**). Within each charge state, there is clear correlation between m/z and cross section and triply charged species split into two prominent subpopulations, as expected from the literature^49–51^. However, the precision of the CCS determination is still more than ten-fold higher than the width of the ion mobility distribution for a given m/z. This results in additional peptide information that can be used for matching and identification.

## DISCUSSION

Here, we have described the construction and evaluated the performance of a state of the art quadrupole time of flight instrument with a trapped ion mobility device and deep integration of the PASEF principle. The novel Bruker *timsTOF Pro* successfully incorporates these building blocks in a robust and flexible manner, not only enabling shotgun-based PASEF operation but many other operation modes, which are still left to be explored.

The full implementation of PASEF in the hard-and firmware in an online format achieved results almost completely in line with those modeled and extrapolated from a laboratory prototype in our 2015 paper^31^. This suggests that the physical operating principles are indeed directly translatable to proteomics workflows. In particular, the instrument routinely delivers sequencing rates above 100 Hz in complex proteome samples. In standard MS/MS acquisition schemes, such high fragmentation rates inevitably imply very short ion collection times and consequently poor spectrum quality. In contrast, PASEF leverages the full scan speed of TOF instruments with undiminished sensitivity as precursor ions are trapped and released as condensed ion packages by the time they are selected for fragmentation. This enabled the identification of over 6,000 protein groups in single runs from a human cancer cell line with minimal input material, and with high quantitative accuracy.

While we focused on label-free quantification in the current study, we expect that the high number of spectra per run will particularly benefit MS/MS-based quantification methods, for example isobaric labeling with TMT^52^, iTRAQ^53^ or EASI-tag^54^. These approaches should additionally benefit from the ion mobility separation itself as it increases the purity of the isolation window and thereby reduces potential artefacts from co-eluting and co-isolated precursor ions.

The high speed and sensitivity of the *timsTOF Pro* allowed us to drastically decrease both measurement time and sample amount, culminated in the identification of about 2,500 proteins from only 10 ng HeLa digest in 30 min. This makes the instrument very attractive for proteomics studies with extremely low starting amounts, for example micro-dissected tumor biopsies, and for high throughput clinical applications of proteomics, in particular in combination with robust and fast LC systems.

Finally, we demonstrated that TIMS-PASEF provides an efficient way to generate comprehensive libraries of peptide collisional cross sections, much beyond past reports^51^. Such large-scale measurements could contribute to elucidating fundamental properties of modified and unmodified peptide ions in the gas phase and may eventually enable the *in silico* prediction of CCS values by deep learning algorithms. Furthermore, the very high precision of the CCS measurements with TIMS demonstrated here opens up new avenues for spectral library-based identifications, in which the CCS parameter adds important evidence either on the MS level or, in data-independent acquisition strategies, also on the MS/MS level.

We conclude that the *timsTOF Pro* is a high performance addition to the technology toolbox in proteomics, with many added opportunities enabled by TIMS-PASEF.

## Acknowledgements

We thank our colleagues in the department of Proteomics and Signal Transduction and at Bruker Bremen and Bruker Billerica for discussion and help, in particular Drs. P. Geyer and I. Paron. This work was partially supported by the German Research Foundation (DFG–Gottfried Wilhelm Leibniz Prize) granted to Matthias Mann and by the Max-Planck Society for the Advancement of Science.

## SUPPLEMENTARY FIGURES

**Supplementary Figure 1.**
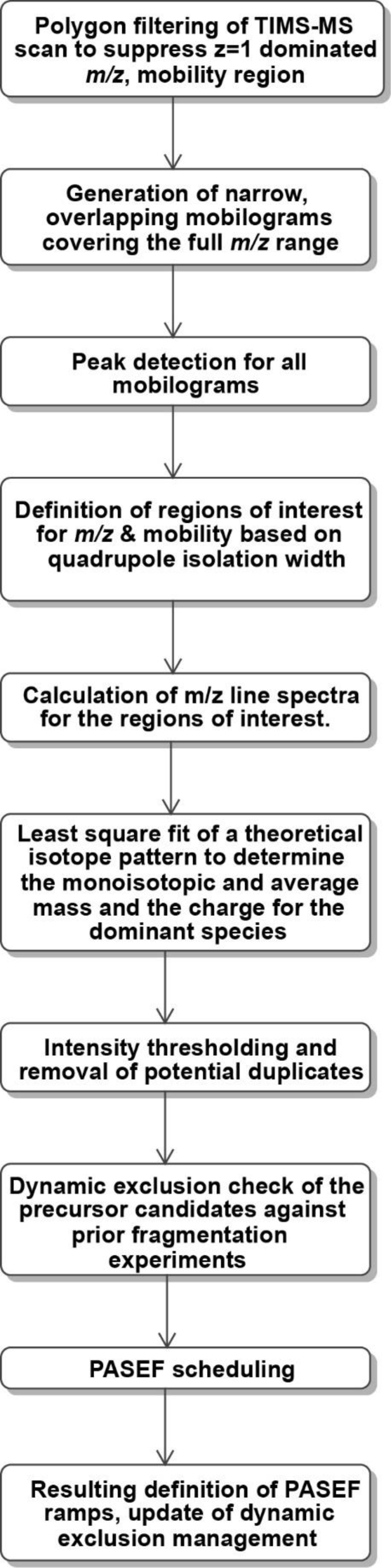
Schematic of the PASEF precursor ion selection scheme.

**Supplementary Figure 2.**
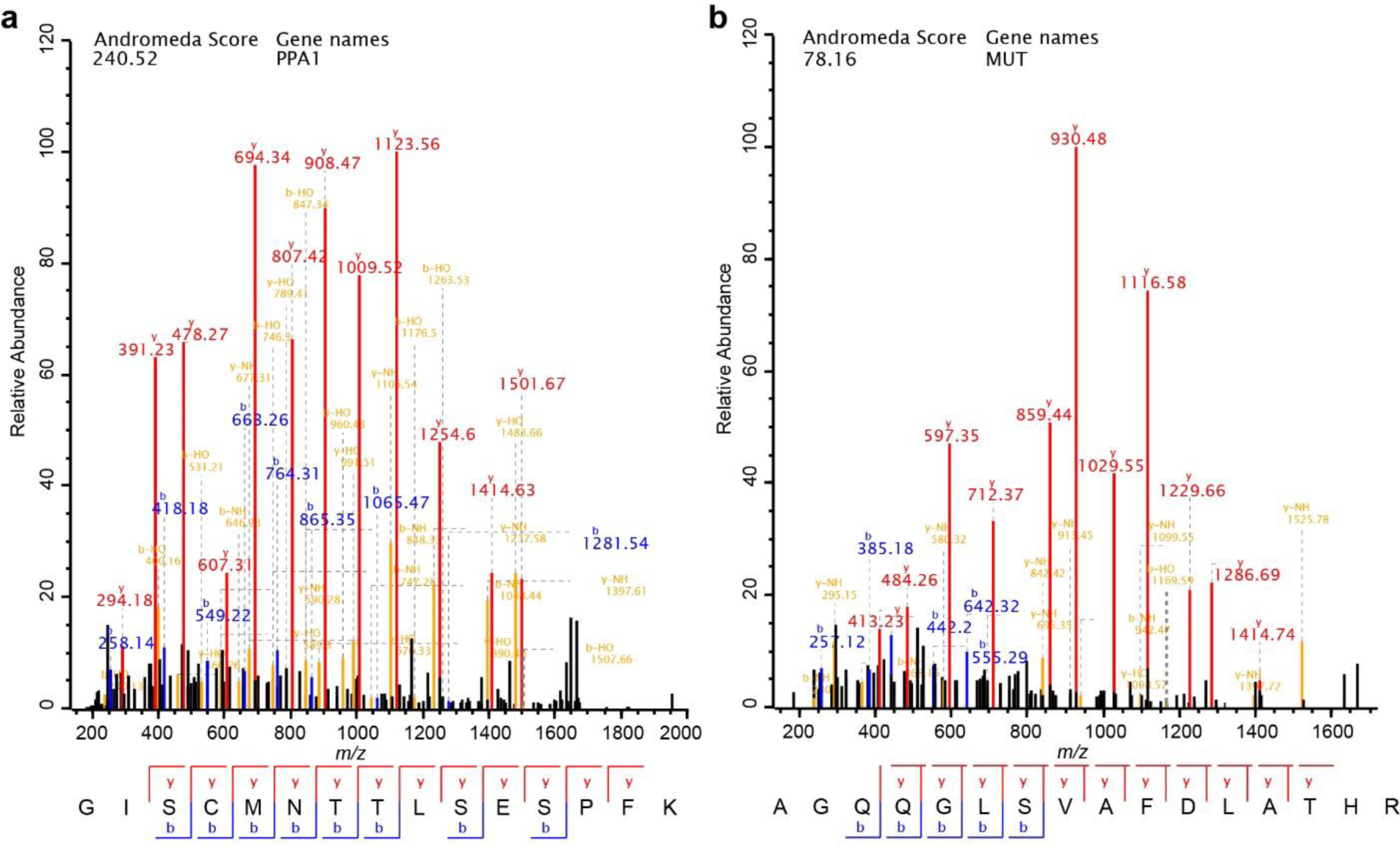
MaxQuant identification of co-eluting peptides of very similar mass, which would have been co-fragmented without TIMS.

